# GlycoPathDB: A database of monosaccharide biosynthesis pathways

**DOI:** 10.1101/2020.10.10.334391

**Authors:** Jaya Srivastava, P. Sunthar, Petety V. Balaji

## Abstract

A distinctive feature of glycans vis-à-vis proteins and nucleic acids is its structural complexity which arises from the huge repertoire of monosaccharides, isomeric linkages and branching. Around 70 monosaccharides have so far been discovered in natural glycans and the pathway for the biosynthesis of 57 of these have been experimentally characterized. However, there is no common platform with a comprehensive information of monosaccharide pathways and enzymes. We have gathered 542 experimentally characterized enzymes of these pathways from literature and set up a first of its kind database called GlycoPathDB (http://www.bio.iitb.ac.in/glycopathdb/). Annotations such as the reaction catalysed, substrate specificity, biosynthesis pathway and PubMed IDs are provided for all the enzymes in the database. Sequence homologs of the experimentally characterized enzymes found in nearly 13,000 completely sequenced genomes from Bacteria and Archaea have also been included in the database. This platform will help in deduction of evolutionary relationships among enzymes such as aminotransferases, nucleotidyltransferases, acetyltransferases and SDR family enzymes. It can also facilitate experimental studies such as direct enzyme assays to validate putative annotations, establish structure-function relationship, expression profiling to determine the function, determine the phenotypic consequences of gene knock-out/knock-in and complementation studies.

## Introduction

Glycosylation is prevalent in all domains of life and is important for a wide range of biological processes. A variety of monosaccharides has been reported to be used by organisms; in particular, those used by prokaryotes seem to be particularly diverse. Such a diverse set of monosaccharides are not available in the growth medium for any of the prokaryotes – they have to be biosynthesized de novo. It is of interest to assess the evolutionary relationships of the biosynthetic pathways and thereby find clues to the need for diversification of glycan alphabet. However, experimental data on the biosynthesis pathways are scattered in literature and hence, it is perceived that organizing this data in a single platform can facilitate studies on evolutionary relationships. Additionally, such a platform can be exploited to identify sequence homologs in whole genome sequences: this can facilitate further experiments such as direct enzyme assays, knock-out/knock-in and complementation studies, expression profiling, etc. Such studies will contribute to establishing structure-function relationships in glycans.

Prokaryotic cell-surface glycans are at the forefront of mediating interactions of the organism with its environment and hence are under constant selection pressure to evolve with varying extracellular conditions. Organisms mediate glycan diversity by either incorporating diverse monosaccharides, or by linkage, branching and chain length variations. The number of monosaccharides found in naturally occurring glycans is very large (70+). Many of these monosaccharides are stereoisomers of each other and are derived from simple sugars (e.g., hexoses) by modifications such as deoxygenation, amination, acetylation, methylation, inversion of configuration, etc. These modifications are introduced by several distinct enzymes belonging to different structural families. Distinctness may also arise in enzymes belonging to the same structural family, which have evolved to define substrate specificity with the help of only a few residues (Figure 1). Variations in specificity can be with respect to the nucleotide moiety (e.g., UDP- and GDP-Glc2NAc 4,6-dehydratase) or the saccharide moiety (e.g., TDP-3-amino-6-deoxy-Glc/Gal acetyltransferase and UDP-Glc/Glc2NAc 6-dehydrogenase).

**Figure 1:**
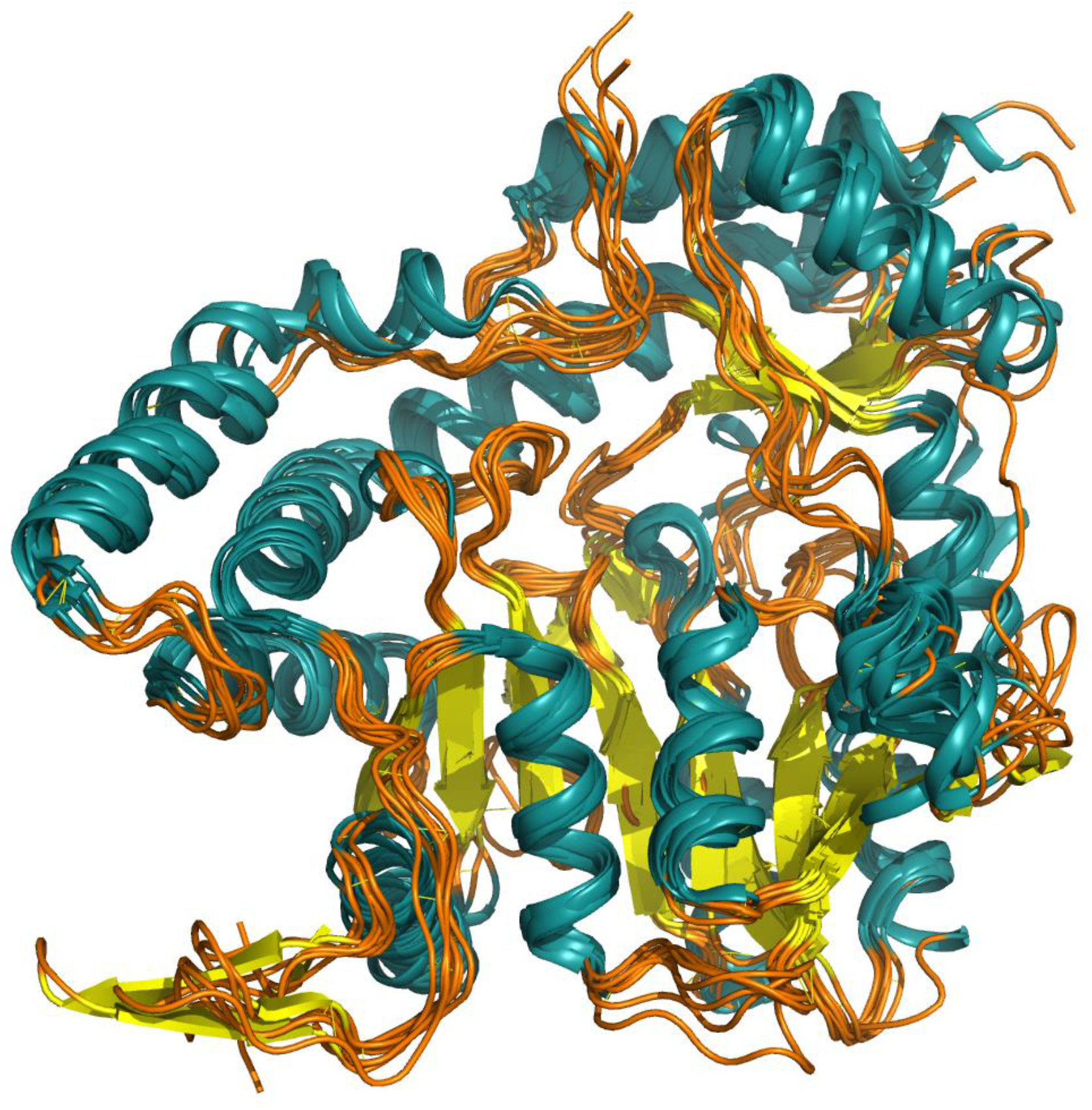
3D structural superimposition of C3- and C4-aminotransferase superfamily enzymes which use different nucleotide sugar substrates. ArnB (PDB id 1MDO, chain A), PseC (2FNI, chain A), DesV (2OGA, chain A), perA (3BN1, chain A), WecE (4PIW, chain A) and PglE (4ZTC, chain A) are C4-aminotransferases, respectively, from the pathways for the biosynthesis of UDP-L-Ara4N, CMP-L-Pse45Ac7Ac, (d)TDP-desosamine, GDP-Per, (d)TDP-Fuc4NAc and CMP-Leg5Ac7Ac. WbpE (3NYU, chain A) and WlarG (5U1Z, chain A) are C3-aminotransferases, respectively, from the pathways for the biosynthesis of UDP-Man2NAc3NAcA and (d)TDP-Fuc3NAc/Qui3NAc.

The mechanism by which modifications in hexoses benefit organisms is underexplored. Information on monosaccharide biosynthesis enzymes is dispersed in literature and a one stop knowledge base is lacking. A common platform which organizes the enormous amount of experimental data on monosaccharide biosynthesis pathways accumulated over the years will facilitate research exploring neo-functionalization and establishing structural and functional themes in these diverse groups of enzymes. It can be exploited in genetic studies aiming to decipher mechanisms by which monosaccharide modifications benefit organisms with varying environmental factors. In this study, we present such an integrated platform which includes 542 experimentally characterized enzymes. Sequence homologs of this curated set of proteins in ~13000 archaeal and bacterial whole genome sequences have also been included in this database. In addition, the approach utilized for functional prediction has been incorporated as a prediction tool on the user interface to check if user-provided sequences are homologous to the monosaccharide biosynthesis enzymes.

## Methods and Implementation

### Overview

Experimental data for monosaccharide biosynthesis enzymes were obtained from literature by keyword search and back references. Based on sequence similarity, enzymes were checked if they are single domain or multi-domain, and domain boundaries were defined in case of the latter. The corresponding UniProt (UniProt Consortium 2019), NCBI (NCBI Resource Coordinators 2018) and PDB (Berman et al. 2000) IDs were fetched. The type of evidence available for the enzyme activity and whether or not the enzyme has been assayed with multiple substrates were checked. Enzymes were assigned functional categories (e.g., dehydratase). Pathways were aggregated based on the precursor and/or nucleotide used for the biosynthesis (e.g., UDP-sugars). Enzymes were grouped into various families based on activity as well as sequence similarity (e.g., C3- and C4-amino transferases). In some cases, the same enzyme could yield more than one product (e.g., transferring -NH2 group in both axial and equatorial orientations). Such enzymes were treated as having broader reaction specificity.

### PUID

Experimentally characterized enzymes are indexed using PUID. It is a unique identifier which consists of UniProt identifier of the protein suffixed with two integers. In case of single domain proteins, the two integers correspond to the residue numbers of N- and C-terminal amino acids. Each domain of a multi-domain protein is stored separately and, in these cases, the two integers correspond to residue numbers of the first and last residue of the domain. This scheme permits unambiguous assignment of functions to corresponding domains in multidomain proteins. For instance, *E. coli* glmU (UniProt id P0ACC7) has a uridylyltransferase domain (residues 1 to 226) and an acetyltransferase domain (residues 227 to 456). These two domains are stored separately with PUIDs P0ACC7_1-226 and P0ACC7_227-456, respectively.

### Data storage and organization

Data are stored in a MySQL database. A relational database is used to accommodate large datasets on the server (Figure 2). Information taken from literature is stored in separate tables. The Experimental Sequence Master table contains protein metadata. Additional metadata are available for all PUIDs (Figure 2). Sequences obtained from UniProt do not contain genome identifiers; hence, these were assigned NCBI genome identifiers of the organism to which sequences belong to. A unique identifier is assigned if the source organism of a UniProt sequence is not recorded in the NCBI database. An enzyme may be associated with one or more pathways depending upon its substrate specificity. This mapping is defined in the Protein Reactions Map table.

**Figure 2:**
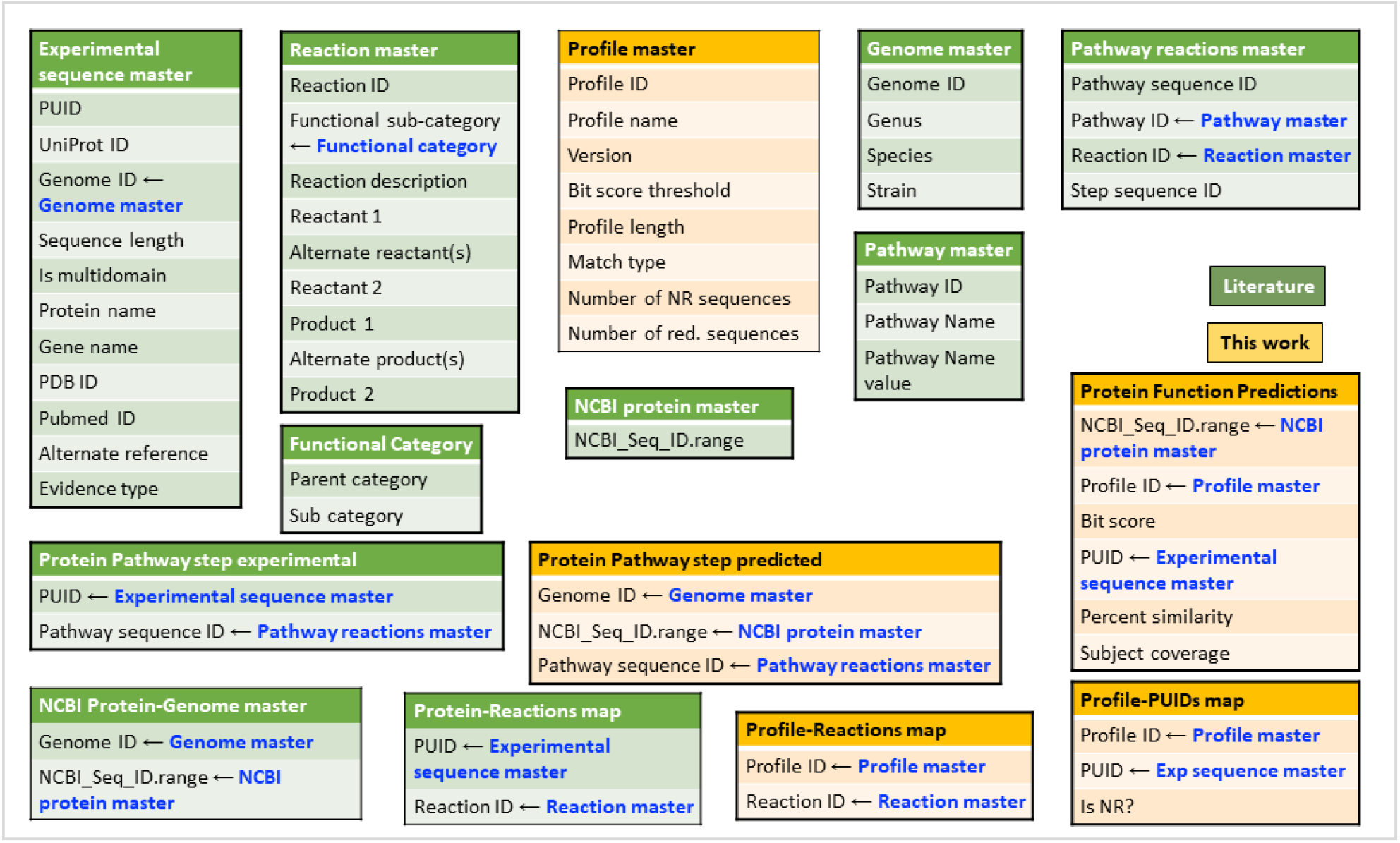
Tables used in GlycoPathDB to store data obtained from literature and manual curation

### Predicted monosaccharide biosynthesis pathways

Experimentally characterized enzymes belonging to monosaccharide biosynthesis pathways were used for sequence homology-based search. For 57 monosaccharides, enzymes catalysing every step of the biosynthesis pathway have been characterized. Most of these enzymes were used to generate HMM profiles (Eddy 2011); the rest were used as BLASTp (Altschul et al. 1990) queries (Srivastava et al, accepted, 10.1099/mgen.0.000452). A HMM profile may be associated with one or more reactions and the Profile Reactions Map table maintains the profile to reaction mapping. These profiles and queries were used to scan whole genome sequences to identify the monosaccharides used by the organism; in all, ~13000 archaeal + bacterial genomes were scanned. These results are stored in Protein Function Predictions and Protein Pathway Step Predicted tables. RefSeq (O’Leary et al. 2016) proteins which were found to be homologs of experimentally characterized proteins of monosaccharide biosynthesis are also stored in the Protein Function Predictions table.

In the NCBI RefSeq database, multiple genomes may contain identical proteins, and hence are referenced by the same identifier. Hence, these identifiers (RefSeq protein IDs) are stored in a separate table called NCBI protein master. They are mapped to respective genomes in NCBI protein-genome master. On the other hand, UniProt contains unique identifiers for sequences which are identical but are from different organisms. Therefore, an additional field for storing genome ID is provided in Protein Pathway step predicted, but is not required for Protein Pathway step experimental. Relationships between these tables is explained in Figure 2. Description of all the fields is documented in supplementary information (Table S1).

### Accessing GlycoPathDB, the web interface

GlycoPathDB is accessible through http://www.bio.iitb.ac.in/glycopathdb/. It is built on the open source Drupal 8 content management system. Composite schematics of the pathways covered in this database as well as individual pathways illustrated schematically are given under the “Monosaccharide biosynthesis pathways” pull down menu; these are static images. The database can be searched using gene name, protein name, UniProt, PDB or PubMed identifier as a keyword using ‘search by keyword’ feature. The database can also be browsed by function, pathway, genome or sequence as described below (Figure 3).

**Figure 3:**
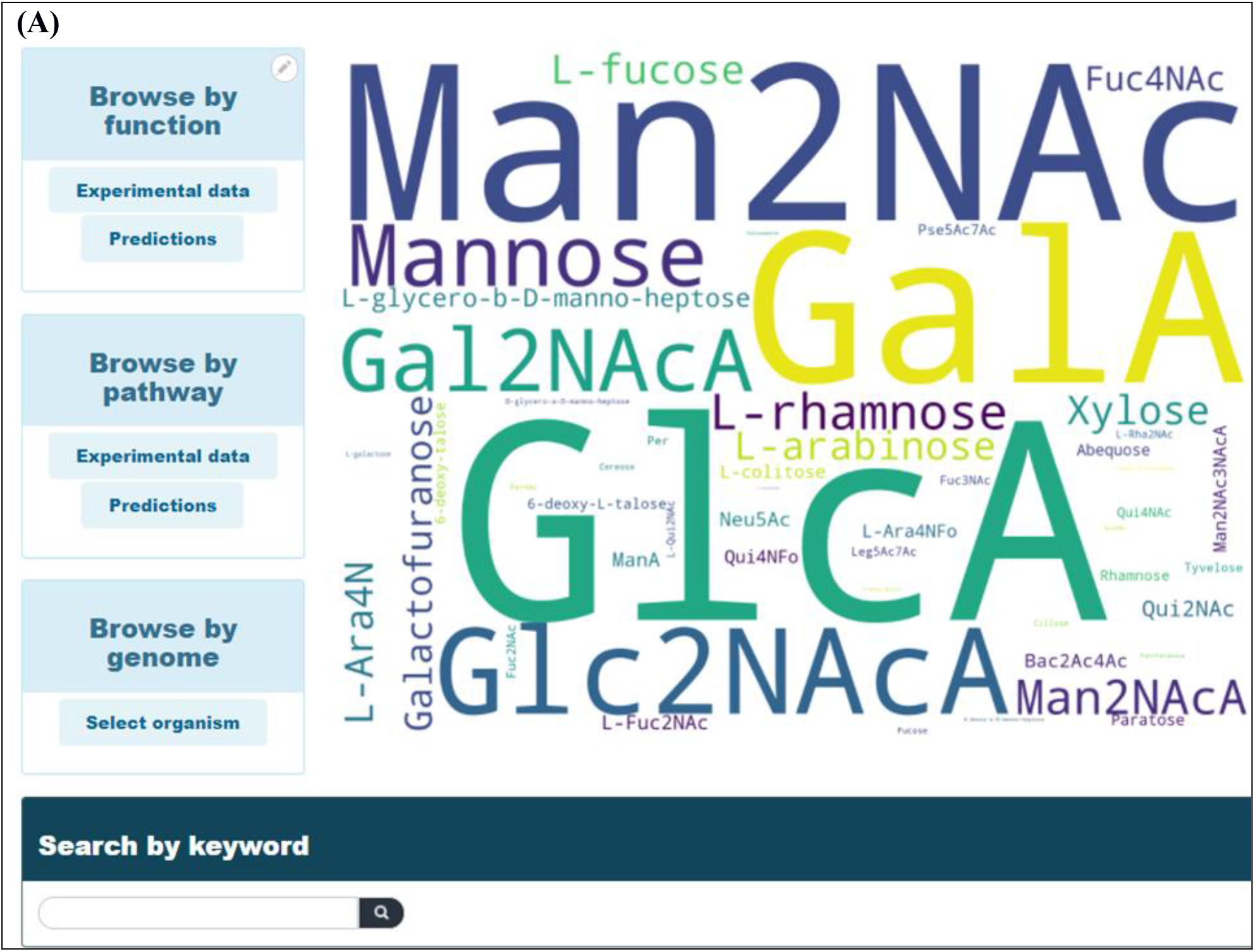
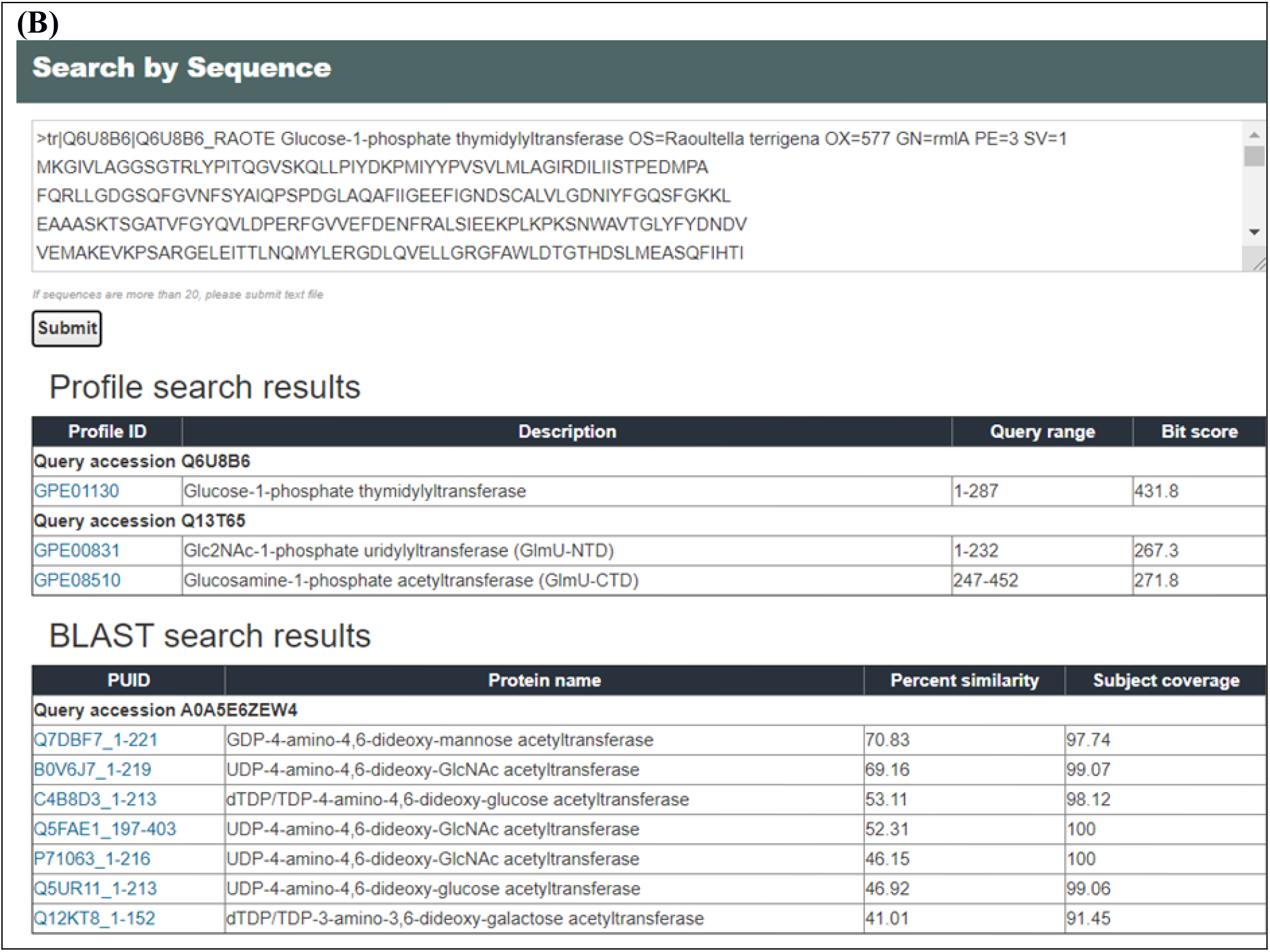
Snapshots of the landing page of GlycoPathDB website. Panel A: Browse features used to search for experimentally characterized and predicted enzymes by function, pathway and by genome. Panel B: Results of search by sequence feature using sample input sequences. Predicted function and corresponding HMM profile or BLASTp scores are displayed.

#### By function

This feature allows a user to search the database for proteins based on a broad level function such as Reductase, Aminotransferase, etc. Where applicable, additional filters are provided to make the search stringent. For example, for the Reductase family, 3-reductase, 4-reductase and reductase are available as three additional filters. Upon selection, PUIDs associated with the selected functional category are displayed. PUID hyperlink displays additional information about the protein e.g., organism, function, length, reaction and PubMed ID of the research article wherein the experimental characterization of enzyme activity is reported.

#### By pathway

This feature can be used to search for organisms which contain a selected pathway from a dropdown menu. Some PUIDs are listed under ‘uncharacterized’ pathway as their sequences were used to generate profiles owing to sequence homology but due to lack of experimental evidence, the biological process/pathway they are involved in is not known.

#### By genome

This feature allows retrieval of monosaccharide biosynthesis pathways encoded by an organism. Drop down menus for genus, species and strain names permit choosing the organism with ease. Results are displayed in two sections: (i) monosaccharides and corresponding experimentally characterized and predicted enzymes of their biosynthesis pathway and (ii) all predictions from the selected genome which participate in monosaccharide biosynthesis.

#### By sequence

Besides retrieving data stored in the database, it is also possible to search the database for homologs of a protein of interest. This is made possible by the ‘search by sequence’ option. HMM profiles and BLASTp queries that were used to scan whole genome sequences have been integrated in the website to find homologs of the sequences input by a user. Results of sample input sequences are illustrated in Figure 3B.

## Results and discussion

Monosaccharide diversity is achieved by enzymes belong to different structural families whose members catalyse other reactions as well. Glucose-1-phosphate, fructose-6-phosphate, sedoheptulose-7-phosphate, UDP-Glc2NAc and GDP-mannose are precursors for the biosynthesis of several monosaccharides (Figure 4). Consequently, they and their derivatives which occur as intermediates are “common” substrates for competing, and functionally diverse, enzymes (Figure 4). Implicit in these assertions is that the end products of the branching pathways are often synthesized by the same organism and at the same stage of growth / life cycle. A particularly illustrative example is that of UDP-Glc2NAc. *Campylobacter jejuni* flagellins are glycosylated with nonulosonates Leg5Ac7Ac and L-Pse5Ac7Ac and their derivatives (Logan et al. 2009). The precursor for both these monosaccharides is UDP-Glc2NAc. This can be acted upon by either a retaining 4,6-dehydratase (Ret) resulting in UDP-4-keto-6-deoxy Glc2NAc (which leads to the biosynthesis of CMP-Leg5Ac7Ac) or an inverting 4,6-dehydratase (Inv) resulting in UDP-2-acetamido-2,6-dideoxy-L-*arabino*-4-hexulose (which leads to the biosynthesis of CMP-L-Pse5Ac7Ac). Here, the terms “retaining” and “inverting” refer to the retention or inversion of the configuration of the pyranose ring C5 atom during the reaction.

**Figure 4:**
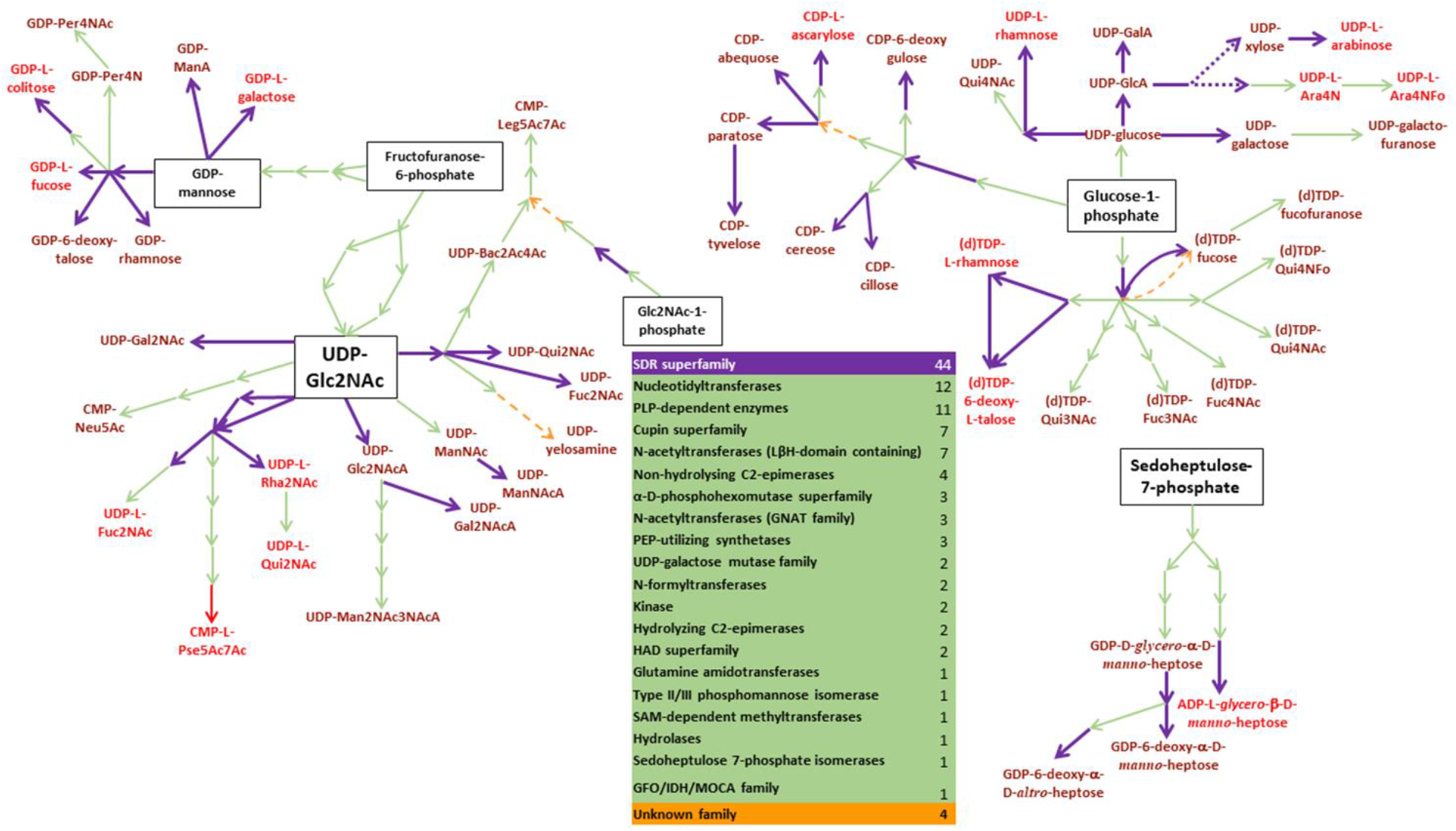
The diversity in monosaccharides arising out of simple hexoses and the enzymes involved in their biosynthesis HAD, haloalkanoic acid dehalogenase Gfo/Idh/MocA, glucose-fructose oxidoreductase/inositol 2-dehydrogenase/rhizopine catabolism protein MocA GNAT, GCN5-related N-acetyltransferases LβH, left handed β helix PEP, phosphoenolpyruvate PLP, pyridoxal 5’-phosphate SAM, S-adenosyl-L-methionine SDR, short chain dehydrogenase reductase

Establishing structure-function relationships among monosaccharide biosynthesis enzymes is a non-trivial task since modification of a few active site residues alters their substrate specificity while preserving the overall structure. For instance, both Inv and Ret are a distinct group of 4,6-dehydratases utilizing UDP-Glc2NAc and share high sequence similarity (>51%). Members of Inv share higher similarity with each other than with those of Ret, and vice versa. Nevertheless, specificity-determining residues are yet to be identified. Counterintuitively, *Bacillus cereus* Pdeg, a retaining dehydratase, is more similar (57-68%) to known inverting dehydratases than to known retaining dehydratases (43-59%). Another example comes from UDP-hexose / hexosamine 4-epimerases which share >45% similarity and exhibit remarkable variations in their substrate specificities viz., UDP-Glc2NAc, -Glc2NAcA, -Glc, -GlcA and -Xyl. While some 4-epimerases such as Gne from *Bacillus subtilis* can utilize both UDP-Glc and UDP-Glc2NAc (Bengoechea et al. 2002), others utilize exclusively only one of these UDP-sugars (Ishiyama et al. 2004). Again, amino acid residues which contribute to specificity in these enzymes have not been characterized. Aside from substrate specificity, functional variations are also observed among homologs: *Escherichia coli* GDP-4-keto-6-deoxy mannose 3-dehydratase (ColD) and *Caulobacter crescenthus* GDP-4-keto-6-deoxy mannose aminotransferase (PerA) share 50% sequence similarity and the same fold (RMSD: 2.7 Å; Figure 5). Two residues which convert the latter (aminotransferase) into the former (dehydratase) have been identified (Cook et al. 2009) but the dehydratase could not be converted into the aminotransferase by reverse mutations.

**Figure 5.**
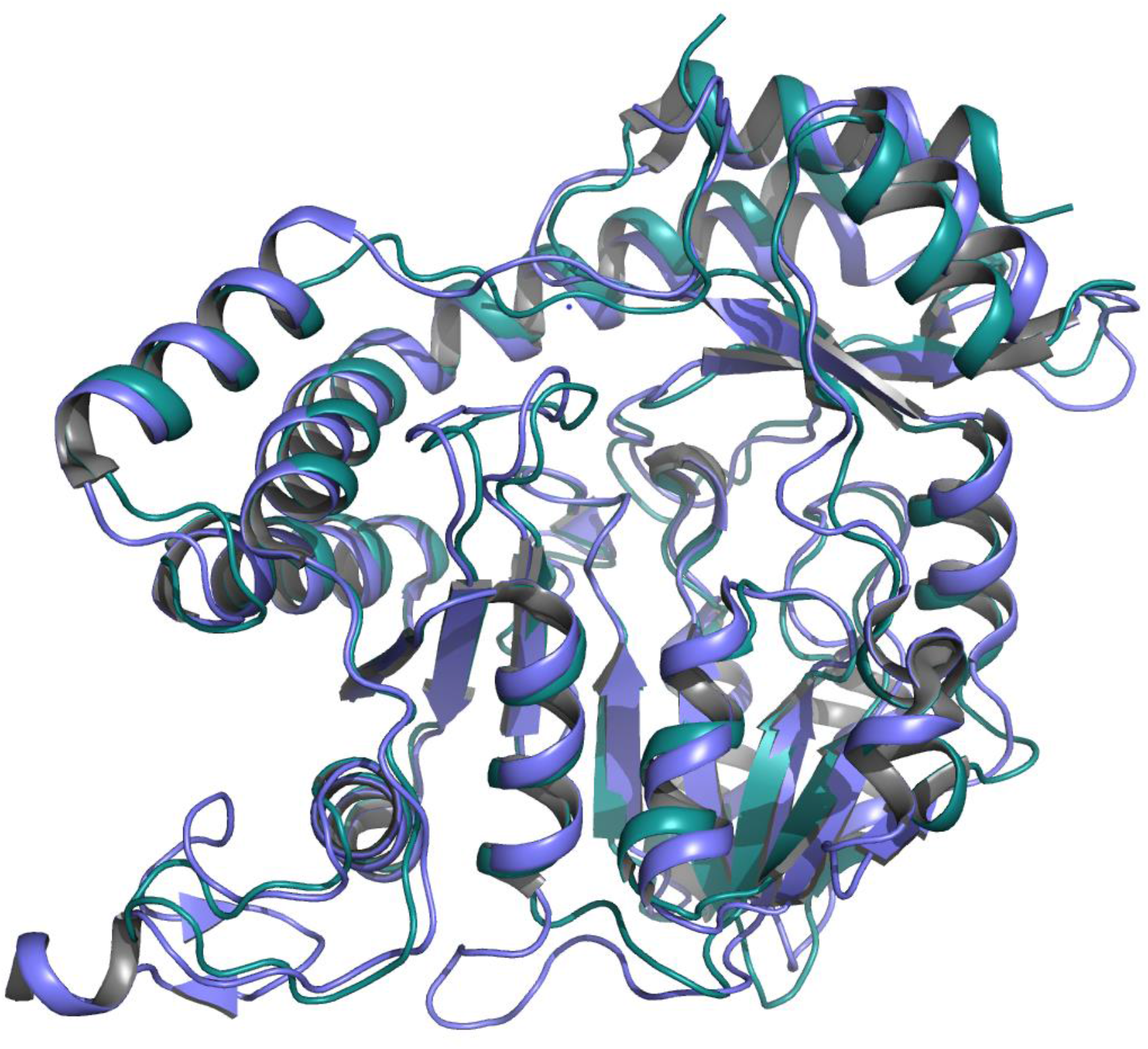
3D structural superimposition of ColD (blue, PDB id 2GMS, chain A), a 3-dehydratase from the GDP-L-colitose biosynthesis pathway, and PerA (teal, 3DR7, chain A), a C4-aminotransferase from the GDP-Perosamine biosynthesis pathway.

The significance of subtle sequence variations within these enzyme families in generating monosaccharide diversity appears to be underappreciated in literature. Figure 4 illustrates a schematic of all monosaccharides whose pathways have been completely characterized in literature and highlights the diversity of participating enzymes. Certain salient features of enzyme families become readily apparent from this composite schematic e.g., a 4,6-dehydratase is the first enzyme in most of the pathways. UDP-Glc2NAc and CMP-Leg5Ac7Ac can be biosynthesised by following two distinctly different pathways. Some epimers (e.g., CDP-abequose and CDP-paratose) are synthesized by enzymes which share significant sequence similarity whereas a few others (e.g., TDP-L-rhamnose and TDP-6-deoxy-L-talose) are synthesized by completely dissimilar enzymes. Aminotransferases and acetyltransferases act in successive steps in the biosynthesis of N-acetyl derivatives; aminotransferases constitute a highly conserved family (figure 1) whereas acetyltransferases belong to two distinct sequence families (table 1). Additionally, those which fall into LbH-domain family contain accessory domains and/or loops for substrate positioning and recognition which are not conserved within all its members (Figure 6). Thus, the ‘why’s and ‘how’s of the evolution of monosaccharide biosynthesis enzymes remain unexplored.

**Table 1:** CATH classification of experimentally characterized N-acetyltransferases involved in the biosynthesis of N-acetyl sugar derivatives and with known 3D structures

| PUID | Gene | PDB ID | Superfamily |  | Fusion protein? |
| --- | --- | --- | --- | --- | --- |
|  |  |  | LbH domain | Domain other than LbH |  |
| LbH domain: present |  |  |  |  |  |
| Q0P9D1_1-195 | PglD | 3BSY | 2.160.10.10 | 3.40.50.20 | No |
| Q6TFC6_1-265 | QdtC | 3FSC | 2.160.10.10 | None | No |
| A9IH93_1-190 | WlbB | 3MQH | 2.160.10.10 | None | No |
| P0ACC7_227-456 | GlmU | 3TWD;1FWY | 2.160.10.10 | 3.90.550.10 | Yes<sup>a</sup> |
| O85353_1-215 | WbqR | 4EA8 | 2.160.10.10 | 3.40.50.20 | No |
| Q12KT8_1-152 | FdtC | 4MZU; 4ZU4 | 2.160.10.10 | 2.60.120.10 | Yes<sup>a</sup> |
| LbH domain: absent |  |  |  |  |  |
| Q8FBQ3_1-224 | WecD | 2FS5 | Not applicable | 3.40.630.30 | No |
| Q383G8_1-147 | GNA1 | 3I3G | Not applicable | 3.40.630.30 | No |
| O25094_1-180 | PseH | 4RI1 | Not applicable | 3.40.630.30 | No |

<sup>a</sup>These proteins have fused domains which perform additional activities

**Figure 6:**
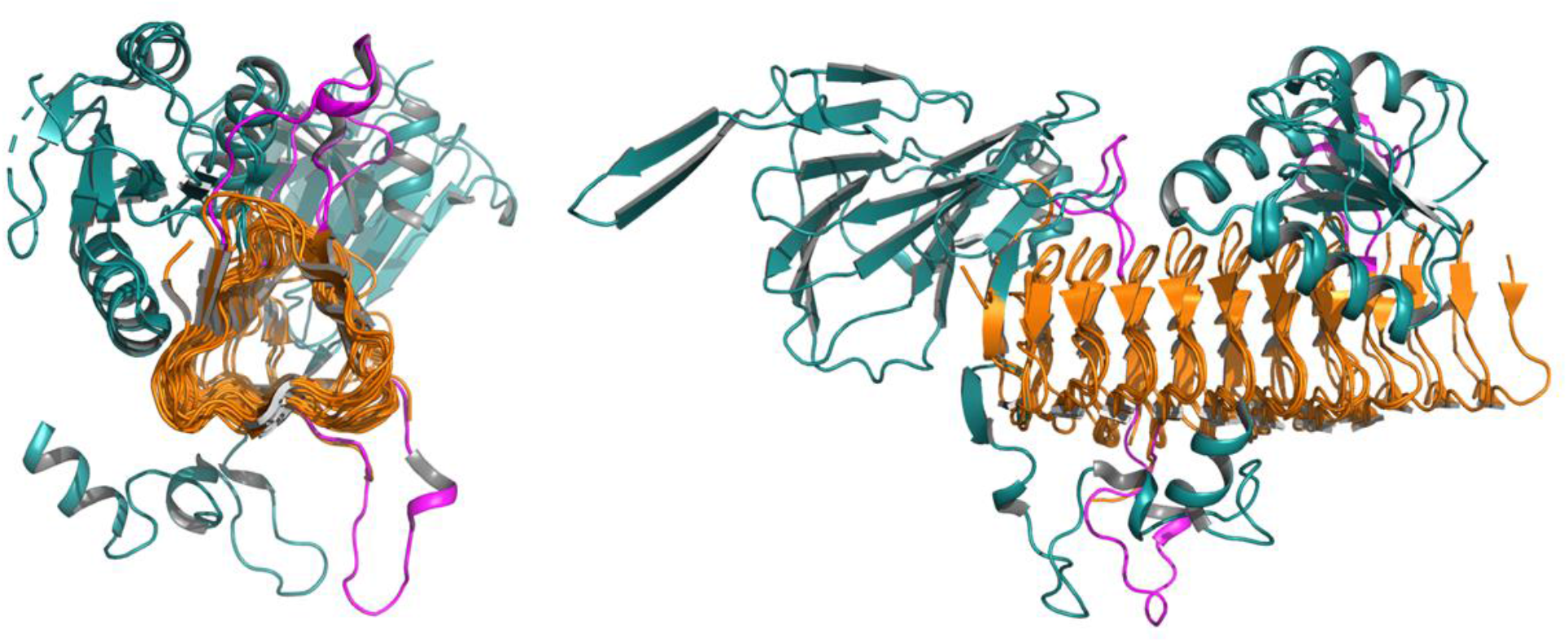
3D structural superimposition of the LbH domain in sugar acetyltransferases viewed along the helix axis (left panel) or perpendicular to the helix axis (right panel). PglD (PDB id 3BSY, chain A), QdtC (3FSC, chain A), WlbB (3MQH, chain A), WbqR (4EA8, chain A) and FdtC (4MZU, chain A) are from the pathways for the biosynthesis of UDP-diNAcBAc, TDP-Qui3NAc, UDP-Man2NAc3NAcA, GDP-Perosamine and UDP-Fuc3NAc, respectively. Functional diversification occurs by the fusion of different domains (rendered in deep teal) at the N- or C-terminus of the LbH domain (in orange) as well as by loop insertions (in pink) within the LbH domain.

Aside from establishing structure-function relationships, these enzymes can therefore be potentially exploited towards evolutionary analysis of origin of diversity in monosaccharides. Glycans are at the forefront of extracellular interactions and monosaccharide diversity is one of the players in creating genus, species, or strain level variations. We believe that a consolidated information of monosaccharide biosynthesis enzymes will benefit these studies.

## Conclusions and future perspective

We have developed a knowledgebase of experimentally characterized enzymes of monosaccharide biosynthesis pathways. Each entry in the database includes gene and protein names, reaction catalysed, substrates utilized, literature reference, enzyme functional category, domain boundaries, source organism, biosynthesis pathway and type of experimental evidence for enzyme activity. The database has hyperlinks to corresponding entries in other databases such as UniProt, PDB and PubMed. Putative sequence homologs of the 542 experimentally characterized enzymes found in 12636 bacterial and 303 archaeal whole genome sequences have also been included in the database. Schematics of all 57 pathways that show ChemDraw^®^ sketches of precursors, intermediates and products are also included. We aim to periodically update the database to cover new reports of monosaccharide biosynthesis pathways (new pathways and existing pathways of different organisms) as well as sequence homologs from other whole genome sequences.

## Acknowledgments

We thank Nitesh Kumar, Ruchi Kumari, Tejas Shah and Tejas Vaidya for technical assistance on the development of GlycoPathDB. Jaya Srivastava is thankful to the Council of Scientific and Industrial Research, Government of India for research fellowship (File number 09/087/(0877)/2017-EMR-I).

## Abbreviations

UDP: Uridine diphosphate
GDP: Guanidine diphosphate
TDP: Thymidine diphosphate
Glc2NAc: 2-N-acetyl-glucosamine
Glc: Glucose
Gal: Galactose
Leg5Ac7Ac: 5,7-N,N’-diacetyl-legionaminic acid
L-Pse5Ac7Ac: 5,7-N,N’-diacetyl-L-pseudaminic acid

