## Supplementary material for "GlycoPathDB: A database of monosaccharide biosynthesis pathways": Table S1

| **Table: Experimental sequence master** | | | | |
| --- | --- | --- | --- | --- |
| **Field** | **Description** | **Field type** | **Data Type** | **Example** |
| PUID | Accession number assigned to a protein. Is of the format UniProt ID_Domain range. | Primary | Text (Plain) | Q9ZGH3_1-337 |
| UniProt ID | Accession number of the protein in the UniProtKB database. | Primary | Text (Plain) | Q9ZGH3 |
| Genome ID | From Genome Master table | Foreign | Same as in Genome Master table | GCF_001443625.1 |
| Sequence length | Number of amino acid residues in the protein. | Primary | Number (integer) | 337 |
| Is multidomain | **Y** for multidomain proteins and **N** for single domain proteins. | Primary | List ([Y,N];Text) | **N** |
| Protein name | Name of the protein as in the UniProtKB database or the NCBI RefSeq database. | Primary | Text (Plain) | dTDP/TDP-glucose 4,6-dehydratase (Retaining) |
| Gene name | Name of the gene that encodes this protein. Taken from the UniProtKB. | Primary | Text (Plain) | desIV |
| PDB ID | Unique identifier in the protein databank for proteins for which 3D structure is known; for others, this will be null. | Primary | Text (Plain) | 1R66;1R6D |
| PubMed ID | Unique integer ID assigned to a record in the PubMed database maintained by the National Library of Medicine, NIH, USA. | Primary | Number (Integer) | 14570895 |
| Alternate reference | URL for research articles for which PubMed ID is not found. | Primary | HTML link | NA |
| Evidence type | Unique ID assigned to denote if the functional annotation of the protein is  1. Inferred from direct enzyme assay (DEA)/Inferred from one pot enzyme assay (OPEA)  3. Inferred from complementation (IC)  4. Inferred from homology (SwissProt) (IH) | Primary | Text (Plain) | DEA/OPEA |

| **Table: Reaction master** | | | | |
| --- | --- | --- | --- | --- |
| **Field** | **Description** | **Field type** | **Data Type** | **Example** |
| Reaction ID | Unique identifier for each enzyme-catalyzed reaction. | Primary | Text (Plain) | 147 |
| Functional sub-category | From Functional category table. Functional sub-category of the enzyme that catalyzes this reaction. | Foreign | Same as in Functional category table | 4,6-dehydratase |
| Reaction description | A description of the reaction. | Primary | Text (Plain) | dTDP/TDP-glucose 4,6-dehydratase (Retaining) |
| Reactant 1 | Substrate (one of the substrates, if there are more than one). | Primary | Text (Plain) | (d)TDP-Glucose |
| Alternate reactant(s) | If the enzyme catalysing this reacting has broad substrate specificity, then substrates other than that listed in the Reactant 1 field are given here. | Primary | Text (Plain) | NA |
| Reactant 2 | Second substrate (applicable only for those reactions which have more than one substrate). | Primary | Text (Plain) | NA |
| Product 1 | Product (major product, if there are more than one product). | Primary | Text (Plain) | (d)TDP-4-keto-6-deoxy-glucose |
| Alternate product(s) | If the enzyme catalysing this reacting has broad substrate specificity, then products formed from ‘alternate reactant’ are given here. | Primary | Text (Plain) | NA |
| Product 2 | Second (minor) product (applicable only for those reactions which have more than one product). | Primary | Text (Plain) | H2O |

| **Table: Profile master** | | | | |
| --- | --- | --- | --- | --- |
| **Field** | **Description** | **Field type** | **Data Type** | **Example** |
| Profile ID | Unique ID assigned to the HMM profile and is of the format GPExxxxx, where xxxxx is a 5-digit number. Each HMM profile is associated with a protein family. | Primary | Text (plain) | GPE05430 |
| Profile name | Name of the profile assigned on the basis of the enzyme activity (or activities) of the proteins belonging to the protein family of this profile. This also is the annotation assigned to a protein which satisfies the bit score threshold of this profile. | Primary | Text (plain) | (d)TDP-glucose 4,6-dehydratase (Retaining) |
| Version | This is used to keep track of the updates to the HMM profile by addition of new sequences that belong to the same protein family and share the same enzyme activity. | Primary | Number (Integer) | 1 |
| Bit score threshold | Best-1-Domain bit score value. A protein sequence which satisfies this threshold is deemed to belong to the same protein family of this HMM profile. | Primary | Number (Integer) | 400 |
| Profile length | Length of the profile | Primary | Number (Integer) | 332 |
| Match type | **Exp** if all the sequences that were used to generate the profile are of type EXP (from the Exp sequence master table); **Extend**, otherwise. | Primary | Text (plain) | **Extend** |
| Number of NR sequences | The number of non-redundant sequences used to generate the profile. | Primary | Number (Integer) | 17 |
| Number of red. sequences | The number of redundant sequences that belong to this protein family. These are the sequences that were not used for generating the profile. | Primary | Number (Integer) | 9 |

| **Table: NCBI protein master** | | | | |
| --- | --- | --- | --- | --- |
| **Field** | **Description** | **Field type** | **Data Type** | **Example** |
| NCBI_Seq_ID.range | NCBI RefSeq id of the protein. Range is of the format *start-end* wherein start and end correspond to the first and last amino acid of the segment of the protein that is identified as a hit to the HMM profile / BLASTp query | Primary | Text (plain) | WP_055643031.1:6-326 |

| **Table: Genome master** | | | | |
| --- | --- | --- | --- | --- |
| **Field** | **Description** | **Field type** | **Data Type** | **Example** |
| Genome ID | From the NCBI RefSeq database. A dummy genome id is assigned if the genome of the organism has not been sequenced. | Primary | Text (plain) | GCF_001443625.1 |
| Genus | Genus to which the organism belongs (from RefSeq database). | Primary | Text (plain) | Streptomyces |
| Species | Species to which the organism belongs (from RefSeq database). | Primary | Text (plain) | venezuelae |
| Strain | Name of the strain (from RefSeq database). | Primary | Text (plain) | ATCC 15439 |

| **Table: Pathway master** | | | | |
| --- | --- | --- | --- | --- |
| **Field** | **Description** | **Field type** | **Data Type** | **Example** |
| Pathway master ID | Accession number assigned to a pathway for the biosynthesis of a monosaccharide. | Primary | Text (plain) | PW042 |
| Pathway name | Name of the biosynthesis pathway. | Primary | Text (plain) | UDP-Gal2NAcA synthesis |
| Pathway name_value* | Name of the biosynthesis pathway | Primary | Text (plain) | UDP-Gal2NAcA synthesis |

*This field is identical to the field Pathway name. It was added to simplify the SQL query

| **Table: Pathway reactions master** | | | | |
| --- | --- | --- | --- | --- |
| **Field** | **Description** | **Field type** | **Data Type** | **Example** |
| Pathway sequence ID | Unique accession number assigned to a step in a biosynthesis pathway | Primary | Text (plain) | 1143 |
| Reaction ID | From Reaction master table. Unique accession number assigned to the step | Foreign | Same as in Reaction master table | 220 |
| Pathway ID | From Path tableway master. The biosynthesis pathway to which this reaction belongs to. | Foreign | Same as in Pathway master table | PW042 |
| Step sequence ID | Serial number; denotes which step of the pathway it corresponds to. | Primary | Number (integer) | 2 |

| **Table: Protein Pathway step experimental** | | | | |
| --- | --- | --- | --- | --- |
| **Field** | **Description** | **Field type** | **Data Type** | **Example** |
| PUID | From Experimental sequence master table. This gives the accession number of an experimentally characterized protein (or a region of the protein) that catalyzes the particular step of the pathway | Foreign | Same as in Experimental sequence master table | Q9ZGH3_1-337 |
| Pathway sequence ID | From Pathway sequence master table. This gives the specific step of the pathway catalyzed by the protein | Foreign | Same as in Pathway sequence master table | 1210 |

| **Table: Protein Pathway step predicted** | | | | |
| --- | --- | --- | --- | --- |
| **Field** | **Description** | **Field type** | **Data Type** | **Example** |
| Genome ID | From Genome master table. This is used to fetch the genus, species and strain names of the organism to which this protein belongs to. | Foreign | Same as in Genome master table | GCF_001443625.1 |
| NCBI_Seq_ID.range | From NCBI Protein master table. This gives the accession number of the protein (or a region of the protein) that catalyzes the particular step of the pathway | Foreign | Same as in NCBI protein master table | WP_055643031.1:6-326;  WP_055643488.1:1-318 |
| Pathway sequence ID | From Pathway sequence master table. This gives the specific step of the pathway catalyzed by the protein | Foreign | Same as in Pathway sequence master table | 1169 |

| **Table: Protein function Predictions** | | | | |
| --- | --- | --- | --- | --- |
| **Field** | **Description** | **Field type** | **Data Type** | **Example** |
| NCBI_Seq_ID.range | From NCBI Protein master table. | Foreign | Same as in NCBI Protein master table | WP_055641634.1:1-327 |
| The following two fields are applicable only if the protein satisfies the bit score threshold of an HMM profile | | | | |
| Profile ID | From Profile master table. The HMM profile to which the protein is a hit i.e., the sequence satisfies the bit score threshold of the profile. | Foreign | Same as in Profile master table | GPE05430 |
| Bit score | Bit score of the sequence. | Primary | Number (integer) | 493 |
| The following three fields are applicable only if the protein is a hit to a BLASTp query | | | | |
| PUID | From Experimental sequence master table. PUID of the BLASTp query. | Foreign | Same as in Experimental sequence master table | NA |
| Percent similarity | Amino acid sequence similarity between the BLASTp query and the hit | Primary | Number (float) | NA |
| Subject coverage | Fraction of the hit protein that is covered in the pairwise alignment with BLASTp query [Note: the query coverage threshold is pre-set for each BLASTp query]. | Primary | Number (integer) | NA |

| **NCBI Protein-Genome master** | | | | |
| --- | --- | --- | --- | --- |
| **Field** | **Description** | **Field type** | **Data Type** | **Example** |
| Genome ID | From Genome master table. This is used to fetch the genus, species and strain names of the organism to which this protein belongs to. | Foreign | Same as in Genome master table | GCF_001443625.1 |
| NCBI_Seq_ID.range | From NCBI protein master table. This gives the accession numbers of proteins contained in this genome | Foreign | Same as in NCBI protein master table | WP_055641634.1:1-327 |

| **Table: Protein-reactions map** | | | | |
| --- | --- | --- | --- | --- |
| **Field** | **Description** | **Field type** | **Data Type** | **Example** |
| PUID | From Experimental sequence master table. Accession number of the protein | Foreign | Same as in Experimental sequence master table | Q9ZGH3_1-337 |
| Reaction ID | From Reaction master table. This is the reaction catalyzed by the protein | Foreign | Same as in Reaction master table | 147 |

| **Table: Profile-reactions map** | | | | |
| --- | --- | --- | --- | --- |
| **Field** | **Description** | **Field type** | **Data Type** | **Example** |
| Profile ID | From Profile master table. Accession number of the HMM profile | Foreign | Same as Profile master table | GPE05430 |
| Reaction ID | From Reaction master table. This is the reaction catalyzed by proteins that were used to generate the profile. | Foreign | Same as Reaction master table | 147 |

| **Table: Profile-PUIDs map** | | | | |
| --- | --- | --- | --- | --- |
| **Field** | **Description** | **Field type** | **Data Type** | **Example** |
| Profile ID | From Profile master table. Accession number of the HMM profile | Foreign | Same as in Profile master table | GPE05430 |
| PUID | From Experimental sequence master table. Accession numbers of proteins that were used to generate the profile | Foreign | Same as in Experimental sequence master table | Q9ZGH3_1-337 |
| Is NR | Indicates if the PUID which is associated with the function described by Profile ID was used to create the profile (Non-redundant) or not (Redundant). Value is **N** for the former and Y for the latter. | Primary | Text (plain) | Y |

| **Table: Functional category** | | | | |
| --- | --- | --- | --- | --- |
| **Field** | **Description** | **Field type** | **Data Type** | **Example** |
| Parent category | Broad level functional category of an enzyme | Primary | Text (plain) | Dehydratase |
| Sub category | Functional sub category | Primary | Text (plain) | (d)TDP-Glucose 4,6-dehydratase |
